# Immediate inhibition of sucrose uptake in *Corynebacterium glutamicum* in response to intracellular glucose-6-phosphate accumulation requires the *ptsG* encoded EII-permease

**DOI:** 10.1101/634246

**Authors:** Dimitar P. Petrov, Oliver Goldbeck, Reinhard Krämer, Gerd M. Seibold

## Abstract

*Corynebacterium glutamicum* co-metabolizes most carbon sources, such as glucose and sucrose. Uptake of those sugars by the PTS involves a glucose- and a sucrose-specific permease EII^Glc^ (*ptsG*) and EII^Suc^ (*ptsS*), respectively. Block of glycolysis by deletion of *pgi* (encodes phosphoglucoisomerase) redirects glucose-driven carbon flux towards pentose phosphate pathway. *C. glutamicum* Δ*pgi* grows poorly with glucose but has unaffected, good growth with sucrose. However, addition of glucose to sucrose-cultivated *C. glutamicum* Δ*pgi* immediately arrested growth via inhibition of the EII^Suc^–mediated sucrose uptake and reduction of *ptsS*-mRNA amounts. Kinetic analyses revealed that sucrose uptake inhibition in *C. glutamicum* Δ*pgi* took place within 15 s after glucose addition. We show that inhibition of PTS-mediated sucrose uptake occurs as direct response to glucose-6-P accumulation. Moreover, addition of non-PTS substrates, which are metabolized to glucose-6-P such as maltose or glucose-6-P itself (uptake was enabled by heterologously produced UhpT), led to similar growth and sucrose uptake inhibition as glucose addition. Despite EII^Glc^ not being involved in uptake of these substrates, negative effects on sucrose uptake after addition of maltose and glucose-6-P were absent in the EII^Glc^–deficient strain *C. glutamicum* Δ*pgi*Δ*ptsG*. These results show that the *ptsG-*encoded EII^Glc^ is part of a novel mechanism for perception of intracellular glucose-6-P accumulation and instantaneous inhibition of EII^Suc^-mediated sucrose uptake in *C. glutamicum*. This novel mode of control of PTS activity by an early glycolytic metabolite probably allows efficient adaptation of sugar uptake to the capacity of the central metabolism during co-metabolization, which is characteristic for *C. glutamicum*.

**IMPORTANCE:** Coordination of substrate uptake and metabolism are a prerequisite for efficient co-utilization of substrates, a trait typical for the Gram-positive *C. glutamicum*. Sucrose uptake via the PTS permease EII^Suc^ in this organism immediately was inhibited in response to intracellular accumulation of the glycolysis intermediate glucose-6-phosphate. This inhibition depends exclusively on the presence but not activity of the PTS permease EII^Gluc^. Thus, *C. glutamicum* possesses a novel, immediate, and PTS-dependent way to control and coordinate both uptake and metabolization of multiple substrates by monitoring of their metabolic levels in the cell. This offers new insights and interesting concepts for a further rational engineering of this industrially important production organism and exemplifies a putative general strategy of bacteria for the coordination of sugar uptake and central metabolism.

## INTRODUCTION

Maintenance of metabolic homeostasis is a crucial regulatory task within microorganisms. Metabolite levels are directly linked to crucial cellular functions such as the control of cell division (1) and the loss of control on internal metabolite levels often results in reduced fitness or non-growth phenotypes (2). Control of metabolic pathways is exerted by a variety of mechanisms on the levels of enzyme activity, transcription, translation, and posttranslational modification (3-5). In that complex regulatory network the control and proper selection of substrate uptake plays a pivotal role (4, 6).

The Gram-positive bacterium *Corynebacterium glutamicum* is widely known for its application in the large-scale industrial production of amino acids (7, 8). It can be cultivated on a variety of single and combined carbon and energy sources (9) and efficiently metabolizes most carbon sources in parallel, *e.g*. the sugars glucose, sucrose, fructose, and maltose (10). With exception of maltose (11), these sugars are taken up and phosphorylated in *C. glutamicum* via substrate specific EII permeases of the PEP: sugar phosphotransferase system (PTS), of which the *ptsG*-encoded EII^Glc^ and the *ptsS*-encoded EII^Suc^ are required for glucose and sucrose uptake, respectively (12) (Fig. 1). Unlike many other bacteria, which show a preference for one substrate above others, in *C. glutamicum* the individual uptake rates of different substrates are well equilibrated to each other during co-utilization (10, 13) and extracellular formation of products indicative for the occurrence of overflow metabolism is not observed for this bacterium under optimal conditions (14). These adaptations require the presence of efficient mechanisms for the coordination of carbohydrate metabolism in *C. glutamicum*.

**Fig. 1:**
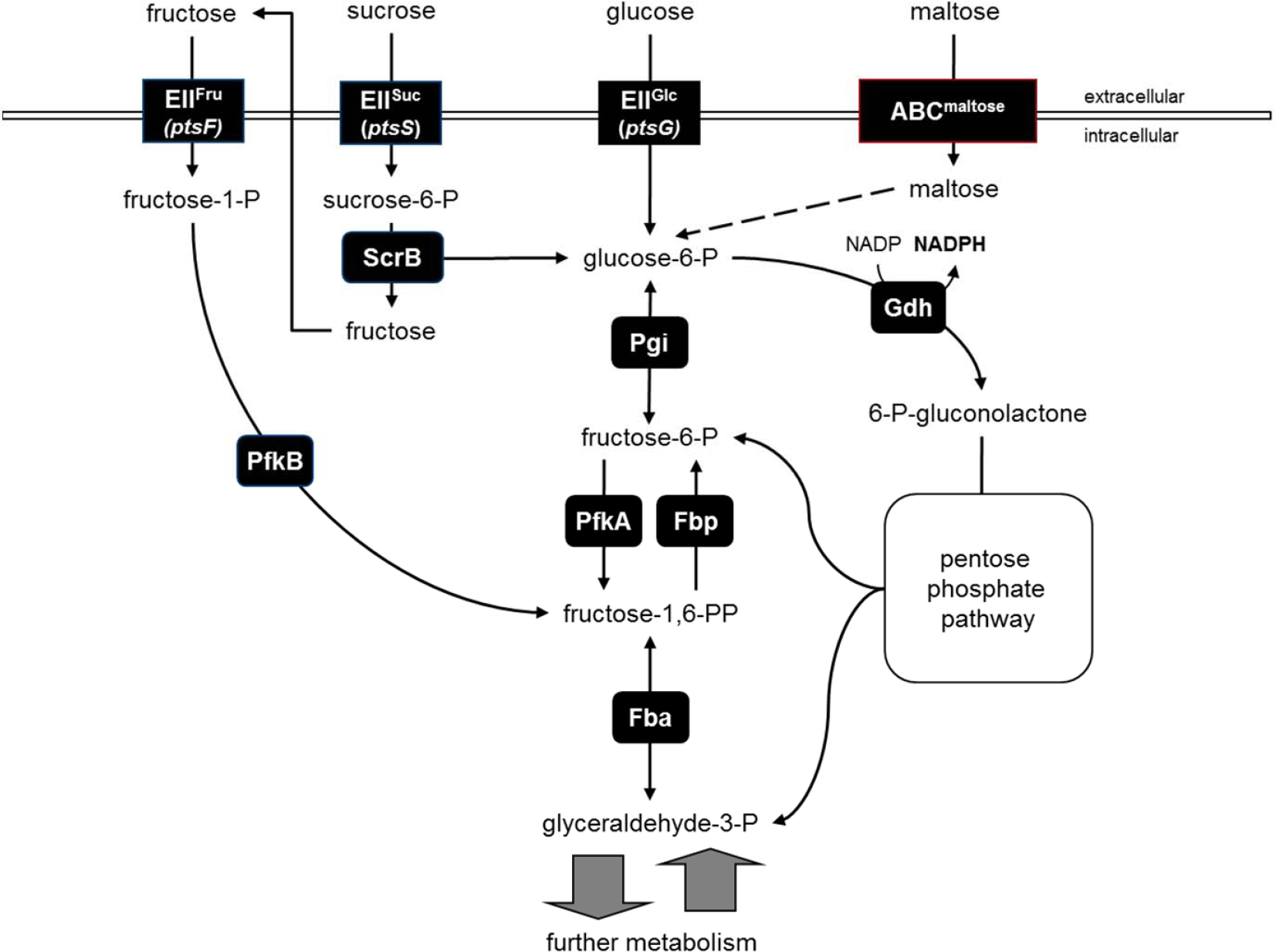
Schematic overview of carbohydrate uptake and metabolism in *C. glutamicum*: Glucose is taken up via the *ptsG* encoded EII^Glc^, thereby formed glucose-6-phosphate is either channeled into glycolysis and then metabolized via 6-phoshofructokinase (PfkA) and fructosebisphosphate aldolase (Fda), or channeled via the glucose-6-phosphate dehydrogenase Gdh into the pentose phosphate pathway. Sucrose is taken up via the *ptsS* encoded EII^Suc^, thereby formed sucrose-6-phosphate is hydrolyzed by the sucrose-6-phosphate hydrolyase ScrB into glucose-6-phosphate and fructose. To phosphorylate thereby formed fructose, fructose is first exported via an unknown transporter and then reimported and thereby phosphorylated by the *ptsF* encoded EII^Fru^. Thereby formed fructose-1-phosphate is phosphorylated by the 1-phosphofructokinase PfkB; resulting fructose-1,6-bisphosphate can be channeled toward the pentose phosphate pathway via the fructose-1,6-bisphosphatase Fbp.

Glucose-6-phosphate (G6P), isomerized by the first step of glycolysis to fructose-6-phosphate (F6P) by the phosphoglucoisomerase (Pgi) (Fig. 1), is an early intermediate of catabolism. G6P is also required for anabolic reactions such as synthesis of cell wall compounds (15). Nevertheless, at high intracellular levels G6P is toxic, impairing cell growth (16, 17) and causing DNA damage (18-20). Therefore, regulatory mechanisms for G6P homeostasis exist: In *E. coli* the small RNA SgrS controls the metabolic stress response that occurs upon accumulation of G6P in *pgi*-deficient strains when cultivated on glucose (21). SgrS binds to *ptsG* transcripts for the glucose specific EIIBC permease of the PTS through Hfq-mediated base pairing at the translation initiation region and thus inhibits *ptsG* translation (22). Moreover, recruitment of RNase E by the SgrS-Hfq ribonucleotide complex leads to fast degradation of the *ptsG* mRNA (23). Similarly, SgrS inhibits also translation of *manXYZ* mRNA thus blocking the production of the broad range PTS sugar transporter ManXYZ (24), which contributes to glucose uptake and phosphorylation (25). In addition, SgrS encodes the small peptide SgrT, which acts as an inhibitor to stop glucose uptake through preexisting EIICB^Glc^ (26, 27). Besides limiting uptake of glucose and concomitant formation of G6P, SgrS mediates also stabilization of the *yigL* mRNA (28). The latter encodes a phosphatase for dephosphorylation of the accumulated G6P forming intracellular glucose (29), which is probably exported via the sugar efflux pumps SetA and SetB (28, 30). Apart from *E. coli* physiological responses to the accumulation of sugar-phosphates have only been scarcely investigated in bacteria including *C. glutamicum.* We recently observed for the Pgi-deficient *C. glutamicum* Δ*pgi* a strong growth defect upon cultivation on glucose, which coincides with drastically reduced glucose-transport activity and *ptsG* transcript amounts (31). This response to the accumulation of G6P in *C. glutamicum* Δ*pgi* resembles the response described for *pgi*-deficent *E. coli* strains (21). However, as no homologues of SgrS, SgrT, or Hfq are present in *C. glutamicum*, different regulatory mechanisms are probably underlying this organisms response towards G6P accumulation.

In this communication we investigated the effects of G6P accumulation in *C. glutamicum* Δ*pgi* on sucrose uptake via the *ptsS* encoded EII^Suc^. We here observed a drastic inhibition of sucrose uptake within seconds after the G6P accumulation was initiated. This inhibition of EII^Suc^ activity by G6P accumulation proceeded independently of the transcriptional and translational control of *ptsS* and exclusively depended on the presence of the *ptsG* encoded EII^Glc^. These results show that beside its function for uptake and concomitant phosphorylation of glucose the *ptsG* encoded EII^Glc^ participates in a novel mechanism underlying the perception of intracellular metabolite accumulation and the fast control of substrate uptake in *C. glutamicum*.

## RESULTS

### Presence of glucose inhibits sucrose utilization in *C. glutamicum* Δ*pgi*

In *E. coli* accumulation of G6P reduces the expression and activity of both uptake systems, which contribute to intracellular formation of G6P - the glucose-specific EIIBC, encoded by *ptsG*, as well as the broad range PTS-sugar transporter ManXYZ (24). *C. glutamicum* does not possess a second PTS permease for glucose uptake (12), but G6P is also formed in the course of sucrose metabolism. As characterized here, sucrose in this bacterium is exclusively taken up via the sucrose-specific, *ptsS*-encoded EII^Suc^ with a V_max_ of 20.4 ± 0.6 nmol*(min*mg dw)^−1^ and a K_m_ of 3.8 ± 0.9 μM (Fig S1). Thereby intracellular sucrose-6-phosphate is formed, which is then hydrolysed by the sucrose-6-phosphate hydrolase ScrB to fructose and G6P (32) (Fig. 1). To study possible effects of intracellular G6P accumulation on *ptsS* transcription and EII^Suc^ activity, the Pgi-deficient strain *C. glutamicum* Δ*pgi* was cultivated in minimal medium with 2% (w/v) sucrose as sole source of carbon and energy. In these cultivations absence of Pgi did not negatively affect utilization of sucrose: Growth, *ptsS*-expression, and sucrose uptake were nearly identical for *C. glutamicum* Δ*pgi* and *C. glutamicum* WT (Fig. 2A); sucrose uptake rates of 19.7 ± 1.8 nmol*(min*mg dw)^−1^ and 19.2 ± 0.9 nmol*(min*mg dw)^−1^ were determined for *C. glutamicum* WT and *C. glutamicum* Δ*pgi*, respectively. In difference to our expectations and unlike the situation for cultivation with glucose, intracellular G6P concentrations were significantly increased in *C. glutamicum* Δ*pgi* when compared to *C. glutamicum* WT in cultivations with sucrose as sole substrate (Fig. S2).

**Fig. 2:**
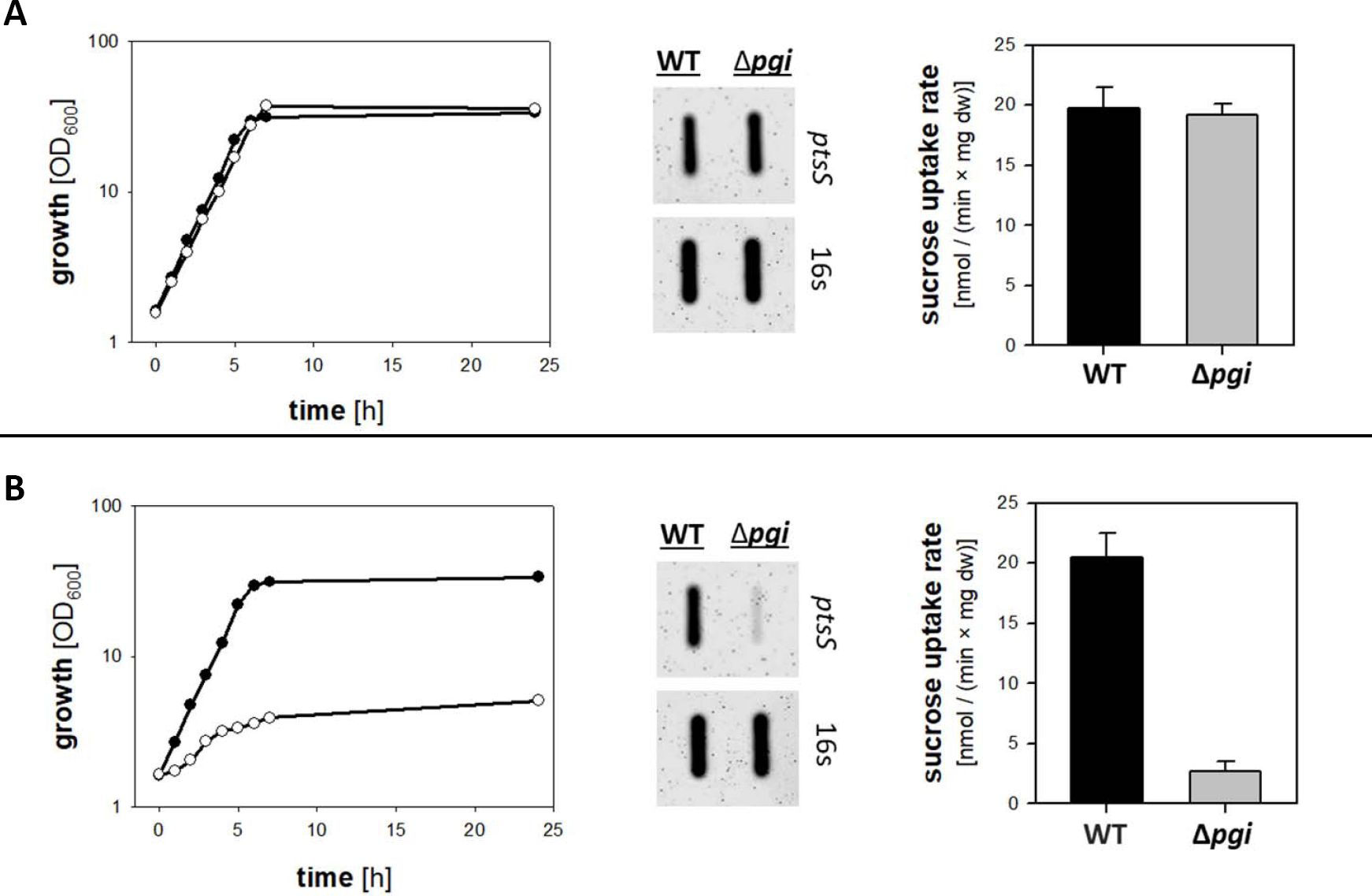
Presence of glucose inhibits sucrose utilization in *C. glutamicum* Δ*pgi*: Growth (right side), *ptsS*-transcript amounts (middle), and [^14^C]-sucrose uptake rates of *C. glutamicum* WT (black circles) and *C. glutamicum* Δ*pgi* (white circles) cells cultivated in minimal medium with 2 % (w/v) sucrose (A) or 1 % (w/v) sucrose plus 1 % (w/v) glucose (B) as carbon and energy source. For growth experiments at least three independent cultivations were performed; data from one representative experiment are shown, results of each of the cultivations were comparable. For analyses of *ptsS* transcript levels in RNA samples from *C. glutamicum* and *C. glutamicum* Δ*pgi* RNA hybridization experiments were performed with specific DIG-labelled antisense RNA probes. Cells for RNA preparation were harvested after reaching an OD_600_ of 6, one representative experiment of a series of three individual experiments is shown. For determination of [^14^C]-sucrose uptake rates of *C. glutamicum* WT and *C. glutamicum* Δ*pgi* cells were harvested at an OD_600_ of 6; data represent mean values and standard deviations of three independent measurements from three independent cultivations.

To investigate effects of internal G6P accumulation on sucrose uptake, we aimed to increase the carbon flux towards G6P formation by cultivation of *C. glutamicum* Δ*pgi* in minimal medium with sucrose plus glucose. Indeed, cultivation of *C. glutamicum* Δ*pgi* with sucrose plus glucose resulted in significantly increased intracellular G6P concentrations when compared to values obtained for *C. glutamicum* WT (Fig. S2). *C. glutamicum* WT grew fast with a rate of 0.42 ± 0.02 h^−1^ in medium with sucrose plus glucose and *ptsS* transcript amounts as well as sucrose uptake activity (20.4 ± 2.8 nmol*(min*mg dw)^−1^) reached about the same levels (Fig. 2B) as determined for cells cultivated with sucrose as sole substrate (Fig. 2A). In difference, growth of *C. glutamicum* Δ*pgi* was significantly slowed down to a rate of 0.09 ± 0.01 h^−1^ and the culture did not exceed an OD_600_ of 4 within 24 h of cultivation when cultivated in minimal medium with sucrose plus glucose (Fig. 2B). Furthermore, the presence of glucose in the culture broth caused for *C. glutamicum* Δ*pgi* a drastic reduction of the *ptsS*-mRNA amounts and significantly decreased sucrose uptake (2.7 ± 0.8 nmol*(min*mg dw)^−1^; Fig. 2B) when compared to *C. glutamicum* WT cultivated with sucrose plus glucose. To rule out that presence of extracellular glucose might inhibit sucrose uptake (e.g. via binding to the permease EII^Suc^), we investigated effects of glucose addition on cultivation of the EII^Glc^ and Pgi-deficient strain *C. glutamicum* Δ*pgi*Δ*ptsG.* Indeed, as shown in Fig. S3, growth, *ptsS* transcription and sucrose uptake were not reduced in cells of *C. glutamicum* Δ*pgi*Δ*ptsG* cultivated with sucrose plus glucose when compared to cells cultivated with sucrose as the sole substrate. Taken together, these results indicate that the EII^Glc^-catalysed uptake and phosphorylation of glucose causes intracellular G6P accumulation in *C. glutamicum* Δ*pgi* cells, which leads to reduced *ptsS* transcript amounts, decelerated EII^Suc^ activity, and slow growth on sucrose plus glucose.

### Transcription initiation of *ptsS* is inhibited in response to intracellular G6P accumulation

To avoid further G6P accumulation upon sugar-P stress, *E. coli* reduces the amounts of *ptsG* and *manXYZ* transcripts by increased transcript degradation rather than lower expression (21, 24). To investigate the mechanisms responsible for the reduced amount of *ptsS* transcript in cells of *C. glutamicum* Δ*pgi* in cultivations with sucrose plus glucose, we determined the *ptsS*-promoter activity using the promoter test vector pET2-PR-*ptsS.* The vector pET2-PR-*ptsS*, carrying the *cat* gene under the control of the *ptsS*-promoter region, was transferred into both *C. glutamicum* and *C. glutamicum* Δ*pgi*. The presence of the plasmid did not change growth of the resulting strains (data not shown). For *C. glutamicum* WT (pET2-PR-*ptsS*) a high *ptsS*-promoter activity was determined for cultivations both on sucrose and on sucrose plus glucose as carbon sources (Fig. 3A). However, whereas the *ptsS*-promoter activity for *C. glutamicum* Δ*pgi* (pET2-PR-*ptsS*) cells cultivated on sucrose as sole carbon source was similar to the values measured for the wild-type reporter strain, the *ptsS*-promoter activity of the *pgi*-deficient cells cultivated with sucrose plus glucose was strongly diminished (Fig. 3A). These results correlated well with the *ptsS*-mRNA levels described above and show that in presence of G6P derived from the added glucose inhibits *ptsS*-expression in *C. glutamicum* Δ*pgi* on the level of transcriptional initiation.

**Fig. 3:**
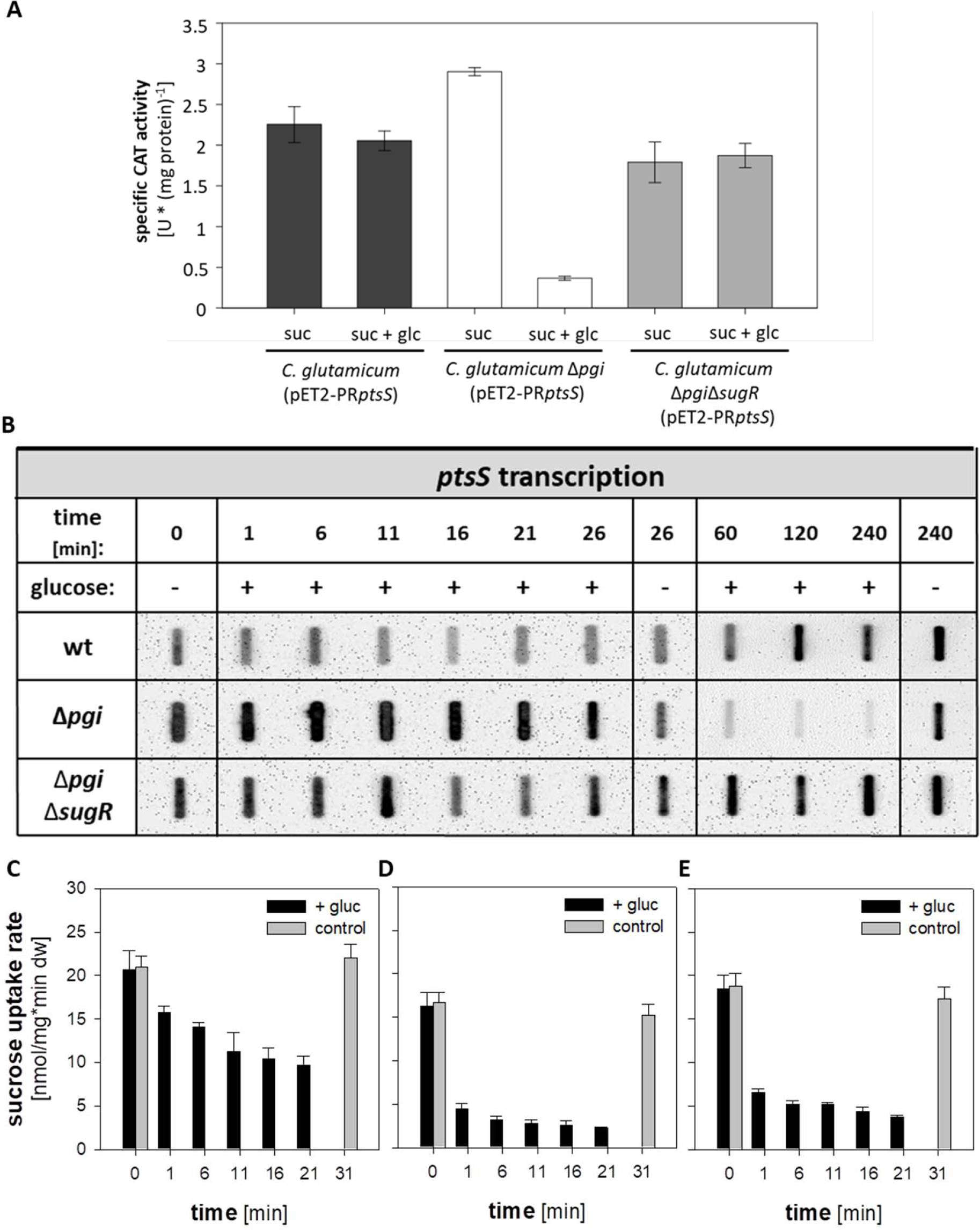
Analysis of *ptsS*-promoter activity in *C. glutamicum* WT (pET-PR*ptsS*) and *C. glutamicum Δpgi* (pET-PR*ptsS*) cells cultivated in minimal medium with 2% (w/v) sucrose or 1% (w/v) sucrose plus 1% (w/v) glucose as substrates (A). Kinetics of the repression *ptsS*-transcription in *C. glutamicum* WT, *C. glutamicum* Δ*pgi*, and *C. glutamicum* Δ*pgi*Δ*sugR* in response to glucose addition (B). Kinetics of the inhibition of [^14^C]-sucrose uptake in *C. glutamicum* WT (C), *C. glutamicum* Δ*pgi* (D), and *C. glutamicum* Δ*pgi*Δ*sugR* (E) cells in response to glucose addition to the culture broth. The reporter gene activity was determined in extracts from cells harvested after 6 hours of cultivation, the values represent averages and standard deviations from at least three independent experiments. For analyses of *ptsS* transcript levels in RNA samples from *C. glutamicum* WT, *C. glutamicum* Δ*pgi*, and *C. glutamicum* Δ*pgi*Δ*sugR* RNA hybridization experiments were performed with a specific DIG-labelled antisense RNA probe. Cells for RNA preparation were harvested at indicated time points before (0 min) and after addition of glucose, one representative experiment of a series of three individual experiments is shown. For determination of [^14^C]-sucrose uptake rates of *C. glutamicum* Δ*pgi* and *C. glutamicum* Δ*pgi*Δ*sugR* cells were sampled at indicated time points and used for transport assays; data represent mean values and standard deviations of three independent measurements from three independent cultivations.

The repressor SugR controls transcription of *ptsS* and further genes coding for PTS components in *C. glutamicum* (33, 34). In the SugR- and Pgi-deficient strain *C. glutamicum* Δ*pgi*Δ*sugR* the presence of glucose during cultivation in minimal medium with sucrose did not lead to reduced amounts of *ptsS* transcripts and lowered *ptsS* promoter activity (Fig. 3A, Fig. S4). These results indicate that SugR-dependent transcriptional control might be responsible for the observed phenotype of *C. glutamicum* Δ*pgi* in cultivations with sucrose plus glucose. However, growth and sucrose uptake of *C. glutamicum* Δ*pgi*Δ*sugR* were slowed down by the presence of glucose in the medium (Fig. S4); the growth rate decreased from a 0.40 ± 0.02 h^−1^ to 0.25 ± 0.03 h^−1^ and the sucrose uptake rate decreased from (2.7 ± 0.8 nmol*(min*mg dw)^−1^to (2.7 ± 0.8 nmol*(min*mg dw)^−1^ for cultivation with sucrose and sucrose plus glucose, respectively. Taken together, these results show that besides the SugR-mediated repression of *ptsS* when *C. glutamicum* Δ*pgi* is cultivated in the presence of glucose, a further regulatory mechanism for the control of sucrose uptake in response to G6P accumulation might be active.

### Sucrose uptake in *C. glutamicum* Δ*pgi* is inhibited instantaneously after glucose addition

To analyze the interplay between the *ptsS*-repression and sucrose uptake inhibition after glucose addition in *C. glutamicum* Δ*pgi*, the dynamics of these two processes was studied in pulse experiments. In detail, *ptsS* transcript amounts as well as sucrose transport rates were determined for cells pre-cultivated with sucrose in regular intervals after the addition of glucose to the culture broth. As expected for *C. glutamicum* WT, the *ptsS* transcript did not decrease within 3 h after glucose addition (Fig. 3B). In accordance, just a slow decrease of sucrose uptake rates from initially 20.6 ± 1.4 nmol*(min*mg dw)^−1^ to 10.4 ± 0.9 nmol*(min*mg dw)^−1^ was observed for *C. glutamicum* WT within 21 min after glucose addition (Fig. 3C). In contrast, the sucrose uptake rate in *C. glutamicum* Δ*pgi* was drastically decreased in the samples taken already 1 min after glucose addition (Fig. 3D). The initial rate of 19.2 ± 0.9 nmol*(min*mg dw)^−1^ dropped to 4.4 ± 0.6 nmol*(min*mg dw)^−1^ and remained low at rates of less than 3.5 nmol*(min*mg dw)^−1^ for the next 25 min of the experiment. Unlike this observed rapid inhibition of sucrose uptake within 1 min, however, the decrease of *ptsS* transcript in *C. glutamicum* Δ*pgi* started only more than 30 min after glucose addition (Fig. 3B). Moreover, also for *C. glutamicum* Δ*pgi*Δ*sugR* we observed a decreased sucrose uptake rate already 1 min after glucose addition (Fig 3E), even though no reduction of *ptsS* transcript was detected after addition of glucose (Fig. 3B). These surprising findings show that the sucrose uptake inhibition occurs prior to reduction of the *ptsS*-mRNA due to the action of SugR and indicates that the inhibition of *ptsS* transcription is not causative for the fast inhibition of sucrose uptake. As expected, no reduction of sucrose uptake was observed when no glucose was added to the culture broth or when glucose was added to the *ptsG*-deficient control-strain *C. glutamicum* Δ*pgi*Δ*ptsG* (Fig. S5).

For a more detailed analysis of the time course of sucrose uptake inhibition by glucose, ^14^C-sucrose uptake experiments with *C. glutamicum* Δ*pgi* cells were pulsed after 45 seconds with unlabeled glucose. The addition of 2 mM glucose to sucrose cultivated *C. glutamicum* Δ*pgi* cells caused an immediate drop of the sucrose uptake rate from initially 19.2 ± 1.3 nmol*(min*mg dw)^−1^ to a rate of 4.5 ± 0.6 nmol*(min*mg dw)^−1^ (Fig. 4A). Extrapolation of the slopes of uptake rates before and after addition of glucose indicated a reaction time for the inhibition of sucrose uptake of less than 1 sec. As in control experiments addition of water instead of glucose did not lead to a decrease of the sucrose uptake rate, it can be ruled out that the effect is due to dilution. In experiments with cells of the EII^Glc^ deficient strain *C. glutamicum* Δ*pgi* Δ*ptsG* the uptake of ^14^C-sucrose was not affected by glucose addition (Fig. 4B). When ^14^C-sucrose uptake was analyzed in pulse experiments with *C. glutamicum* WT a much less pronounced, but significant drop of the transport rate from initially 19.1 ± 1.7 nmol*(min*mg dw)^−1^ to 13.8 ± 0.8 nmol*(min*mg dw)^−1^ was observed immediately after glucose addition (Fig. 4C). These results show that the inhibition of sucrose uptake observed for *C. glutamicum* Δ*pgi* and *C. glutamicum* WT is not caused by competitive inhibition of EII^Suc^ by extracellular glucose. The significantly stronger inhibition observed for the Pgi-deficient *C. glutamicum* Δ*pgi* when compared to *C. glutamicum* WT indicates that a limited capacity for G6P utilization probably is the reason for the observed fast and strong inhibition of EII^Suc^.

**Fig. 4:**
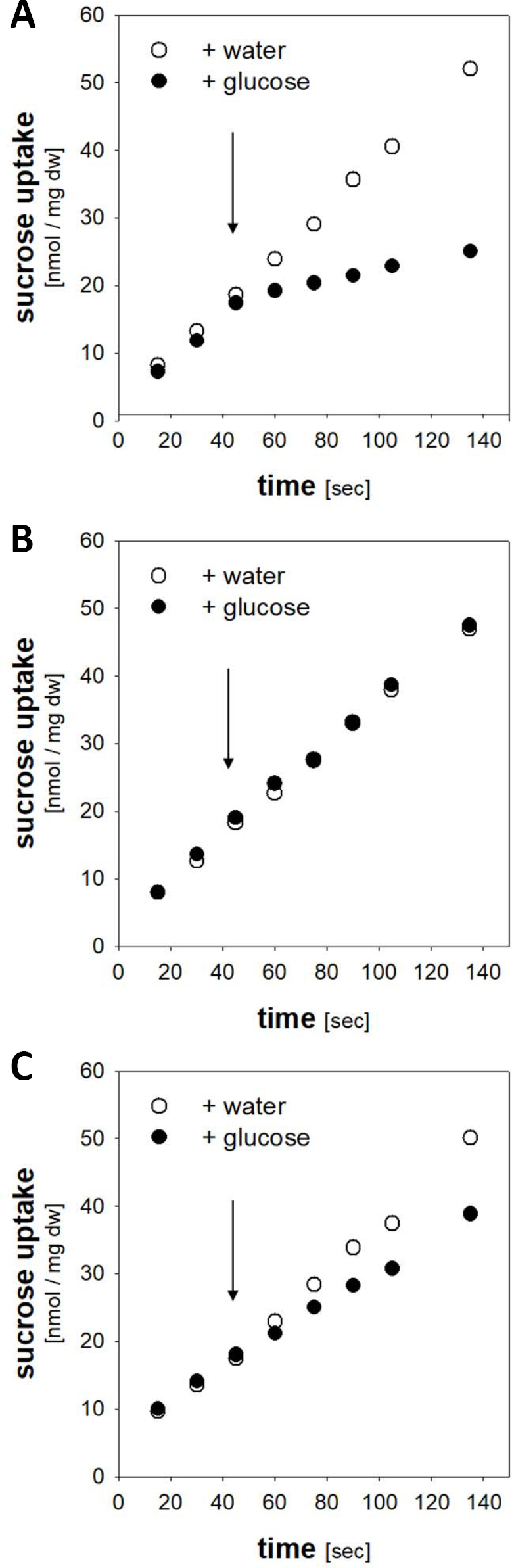
Kinetics of [^14^C]-sucrose uptake in response to glucose addition in cells of *C. glutamicum* Δ*pgi* (A), *C. glutamicum* WT (B), and *C. glutamicum* Δ*pgi* Δ*ptsG* (C). At the time point indicated by the arrow either 70 μl of 200 mM glucose solution (black circles) or 70 μl of water (white circles) were added. One representative experiment of a series of at least three individual experiments is shown.

The instant response of sucrose uptake inhibition observed for *C. glutamicum* Δ*pgi* cells upon addition of glucose makes the involvement of transcriptional and translational regulatory processes unlikely. Indeed, the inhibition of sucrose uptake by glucose occurred to an identical extent when *C. glutamicum* Δ*pgi* cells were pretreated with transcriptional (rifampicin) or translational (chloramphenicol) inhibitors (Fig. S6), when compared to the aforementioned experiments with non-treated cells (Fig. 4). Hence, these results demonstrate that the sucrose uptake inhibition in *C. glutamicum Δpgi* occurred within at least 15 sec after glucose addition and thus far ahead of *ptsS* transcriptional repression which means that a transcription and translation independent regulatory process is involved.

### Fast sucrose uptake inhibition in *C. glutamicum* Δ*pgi* is triggered by intracellular G6P accumulation

The PTS permeases EII^Suc^ and EII^Glc^ depend for their activity on the same PTS phosphorelay mediated by the common components HPr and EI. The strong inhibition of sucrose uptake in the presence of glucose in *C. glutamicum* Δ*pgi* could be caused by two events: (i) a specific stress response caused by the accumulation of G6P or (ii) an insufficient supply of phosphoryl groups for the two permeases EII^Suc^ and EII^Glc^ via the common PTS components EI and HPr. To analyze if PEP limitation or a G6P stress response mechanism is the underlying reason for the observed rapid inhibition of sucrose uptake in *C. glutamicum* Δ*pgi* after glucose addition it is required to uncouple intracellular G6P formation from PTS mediated sucrose uptake. Maltose is taken up in *C. glutamicum* up via the ABC transporter MusEFGK_2_I and metabolized to G6P without concomitant PEP consumption (11, 35). As expected, analyses of intracellular G6P concentration showed that presence of maltose in the culture broth caused significantly increased intracellular G6P concentrations in *C. glutamicum* Δ*pgi* when compared to *C. glutamicum* WT (Fig. S7). This G6P accumulation coincidences with a slowed down growth in medium with 1 % (w/v) sucrose plus 1 % (w/v) maltose of *C. glutamicum* Δ*pgi* [growth rate 0.29 ± 0.01 h^−1^] when compared to *C. glutamicum* WT [growth rate 0.46 ± 0.02 h^−1^] (Fig. 5A). In pulse experiments addition of unlabeled maltose caused a similarly rapid inhibition of ^14^C-sucrose uptake in *C. glutamicum* Δ*pgi* (Fig. 5B) as above described for glucose: Within 15 sec after maltose addition the sucrose uptake rate of *C. glutamicum* Δ*pgi* dropped from initially 19.2 ± 0.9 nmol/(min*mg dw)^−1^ to 8.6 ± 0.6 nmol/(min*mg dw)^−1^. However, growth on maltose as a sole substrate is also slowed down in *C. glutamicum* Δ*pgi* when compared to *C. glutamicum* WT (Fig. S8), indicating that maltose uptake via MusEFGK_2_I itself is subject to G6P accumulation dependent regulation.

**Fig. 5:**
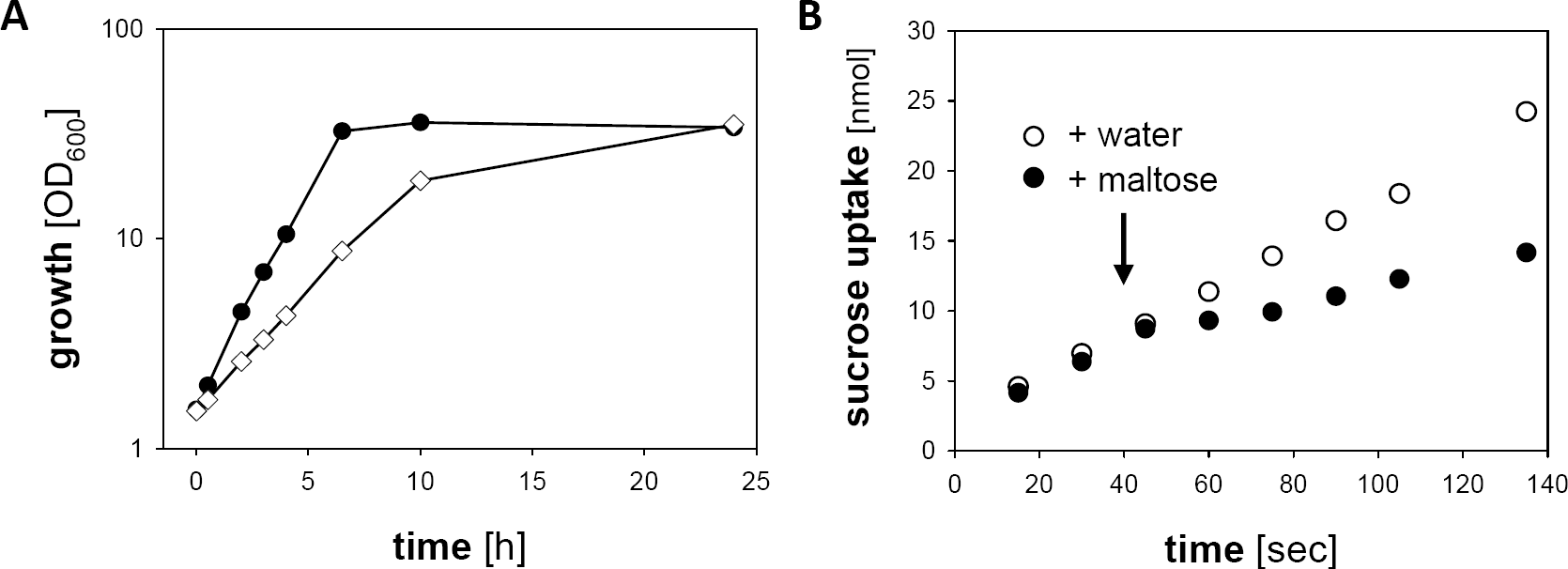
Growth of *C. glutamicum* WT (black circles) and *C. glutamicum* Δ*pgi* (white squares) in CgC minimal medium with 1 % (w/v) sucrose plus 1 % (w/v) maltose as carbon sources [A]; data from one representative cultivation experiment are shown, results of each of the cultivations were comparable. Kinetics of [^14^C]-sucrose uptake in response to maltose addition in cells of *C. glutamicum* Δ*pgi* [B]; at the time point indicated by the arrow either 70 μl of 200 mM glucose solution (black circles) or 70 μl of water (white circles) were added. One representative experiment of a series of at least three individual experiments is shown.

To further evaluate the role of EII^Glc^ for the control of EII^Suc^ activity, we introduced UhpT from *E. coli* as heterologous transporter in *C. glutamicum.* UhpT mediates uptake of G6P in exchange for inorganic phosphate (36, 37). Heterologous expression of *E. coli uhpT* using the plasmid pEKEx2-*uhpT* enabled growth of the resulting strains *C. glutamicum* (pEKEx2-*uhpT*) and *C. glutamicum* Δ*pgi* (pEKEx2-*uhpT*) with G6P as sole source of carbon and energy (growth rates of 0.18 ± 0.04 h^−1^ and 0.19 ± 0.03 h^−1^, respectively) (Fig. S9A and S9B). No growth on G6P was observed for *C. glutamicum* WT (data not shown) and the empty-plasmid control strains *C. glutamicum* (pEKEx2) and *C. glutamicum* Δ*pgi* (pEKEx2) [Fig. S9A and S9B]. For uptake of ^14^C-labelled G6P in *C. glutamicum* (pEKEx2-*uhpT*) a K_m_ of 215 ± 26 μM and a V_max_ of 15.2 ± 0.5 nmol * (min * mg dw)^−1^ were determined. A similar uptake rate of 17.4 ± 1.1 nmol * (min * mg dw)^−1^ and a K_m_ of 150 ± 34 μM for G6P were determined for *C. glutamicum* Δ*pgi* (pEKEx2-*uhpT*) [Fig. S9C]. ^14^C-sucrose uptake experiments with pulses of unlabelled G6P were performed to analyse inhibition of EII^Suc^. The addition of G6P to *C. glutamicum* Δ*pgi* (pEKEx2−*uhpT*) cells caused a rapid inhibition of sucrose uptake from initially 19.5 ± 1.4 nmol * (min * mg dw)^−1^ to 4.8 ± 0.9 nmol * (min * mg dw)^−1^ (Fig. 6A). Also growth of *C. glutamicum* Δ*pgi* (pEKEx2−*uhpT*) on sucrose was slowed down in presence of G6P in the culture broth with growth rates of 0.42 ± 0.02 h^−1^ for cultivation on sucrose and 0.20 ± 0.02 h^−1^ on sucrose plus G6P (Fig. 6B). Growth of *C. glutamicum* Δ*pgi* (pEKEx2-*uhpT*) on sucrose plus G6P proceeds similar as growth with G6P as sole substrate [0.22 ± 0.02 h^−1^]. As expected, the presence of G6P had no effect on sucrose uptake or growth on sucrose for the control strain *C. glutamicum* Δ*pgi* (pEKEx2) [Fig. 6 C & D], which neither grows on G6P nor takes up G6P. Taken together, these results show that PTS-mediated sucrose uptake is also rapidly reduced in *C. glutamicum* Δ*pgi* by the addition of a carbon source such as maltose, which leads to intracellular formation of G6P, but does not require PEP consumption for its uptake.

**Fig. 6:**
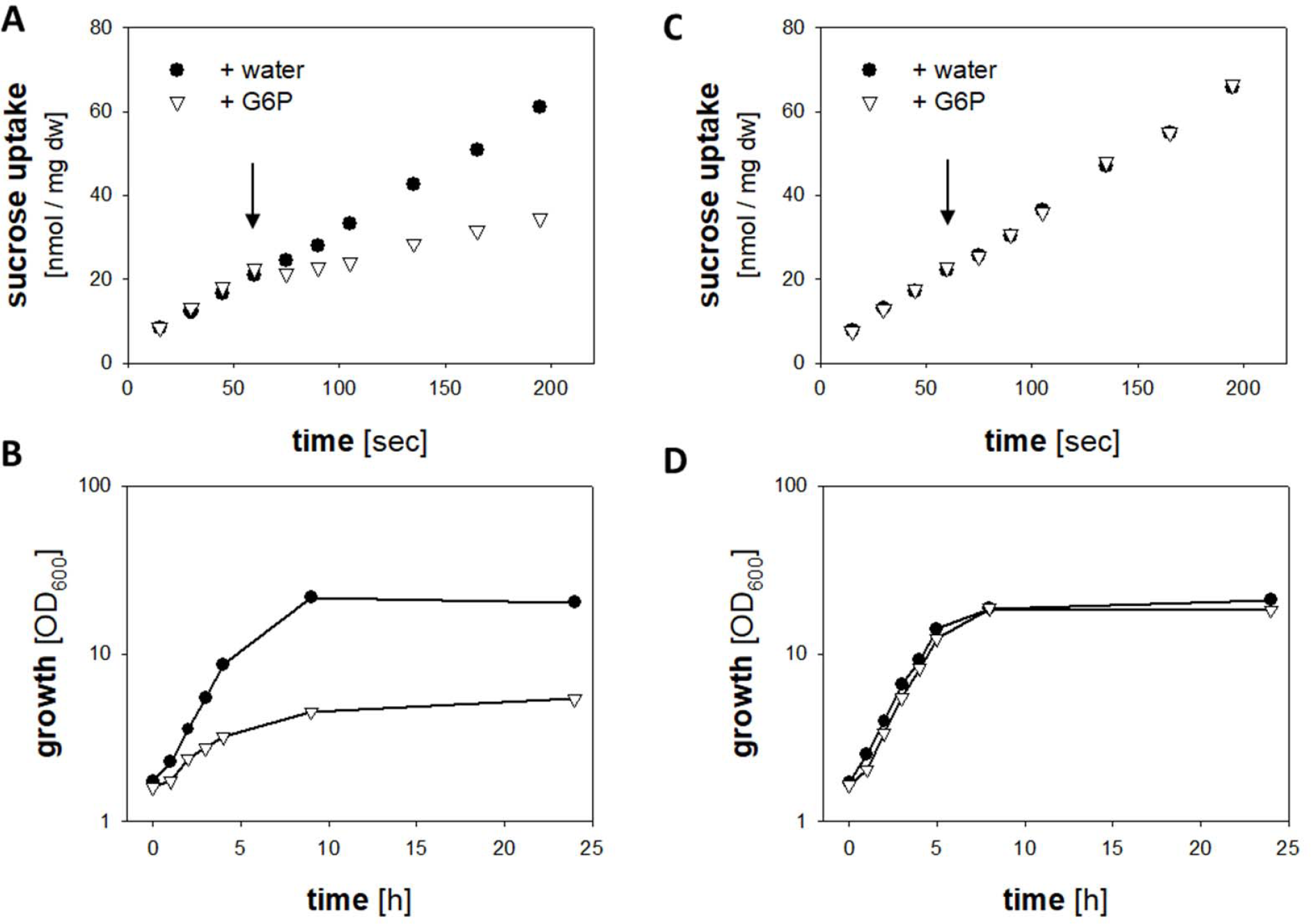
Kinetics of [^14^C]-sucrose uptake in response to maltose addition in cells of *C. glutamicum* Δ*pgi* (pEKEx2-*uhpT*) [A] and *C. glutamicum* Δ*pgi* (pEKEx2) [C]; at the time point indicated by the arrow either 70 μl of 200 mM G6P solution (white triangles) or 70 μl of water (black circles) were added, one representative experiment of a series of at least three individual experiments is shown. Growth of *C. glutamicum* Δ*pgi* (pEKEx2-*uhpT*) [B] and *C. glutamicum* Δ*pgi* (pEKEx2) [D] in CgC minimal medium with 2% (w/v) sucrose [black circles] or 1% (w/v) sucrose plus 1 % (w/v) G6P [white triangles] as sole carbon source, one representative experiment of a series of three individual experiments is shown.

### Fast inhibition of sucrose uptake in *C. glutamicum* Δ*pgi* by intracellular G6P accumulation requires EII^**Glc**^

The immediate onset of sucrose uptake inhibition observed for *C. glutamicum* Δ*pgi* cells upon addition of glucose, maltose, or G6P might either originate from the direct inhibition of EII^Suc^ by intracellular G6P or might be due to a short regulatory cascade directly involving a PTS component. To address that question we tested effects of maltose addition on growth and sucrose uptake inhibition in the EII^Glc^-deficient strain *C. glutamicum* Δ*pgi*Δ*ptsG.* Intriguingly, growth of *C. glutamicum* Δ*pgi*Δ*ptsG* in minimal medium with sucrose was not slowed down by addition of maltose (Fig. 7A), albeit the strain intracellularly accumulated high levels of G6P (Fig. S7). Moreover, no inhibition of ^14^C-sucrose uptake was observed for *C. glutamicum* Δ*pgi*Δ*ptsG* upon addition of unlabelled maltose in pulse experiments (Fig. 7B). Upon ectopic expression of *ptsG* using plasmid pBB1-*ptsG*, in pulse experiments the inhibition of ^14^C-sucrose uptake by maltose addition was restored with *C. glutamicum* Δ*pgi*Δ*ptsG* (pBB1-*ptsG*) to the same level as it was also determined by glucose addition to this strain (Fig. S10). As expected, no inhibition of ^14^C sucrose uptake by maltose addition was observed for the empty-plasmid control strain *C. glutamicum* Δ*pgi*Δ*ptsG* (pBB1) (Fig. S10). These results indicate that EII^Glc^ is required for the perception of G6P accumulation, which is required for the inhibition of EII^Suc^.

**Fig. 7:**
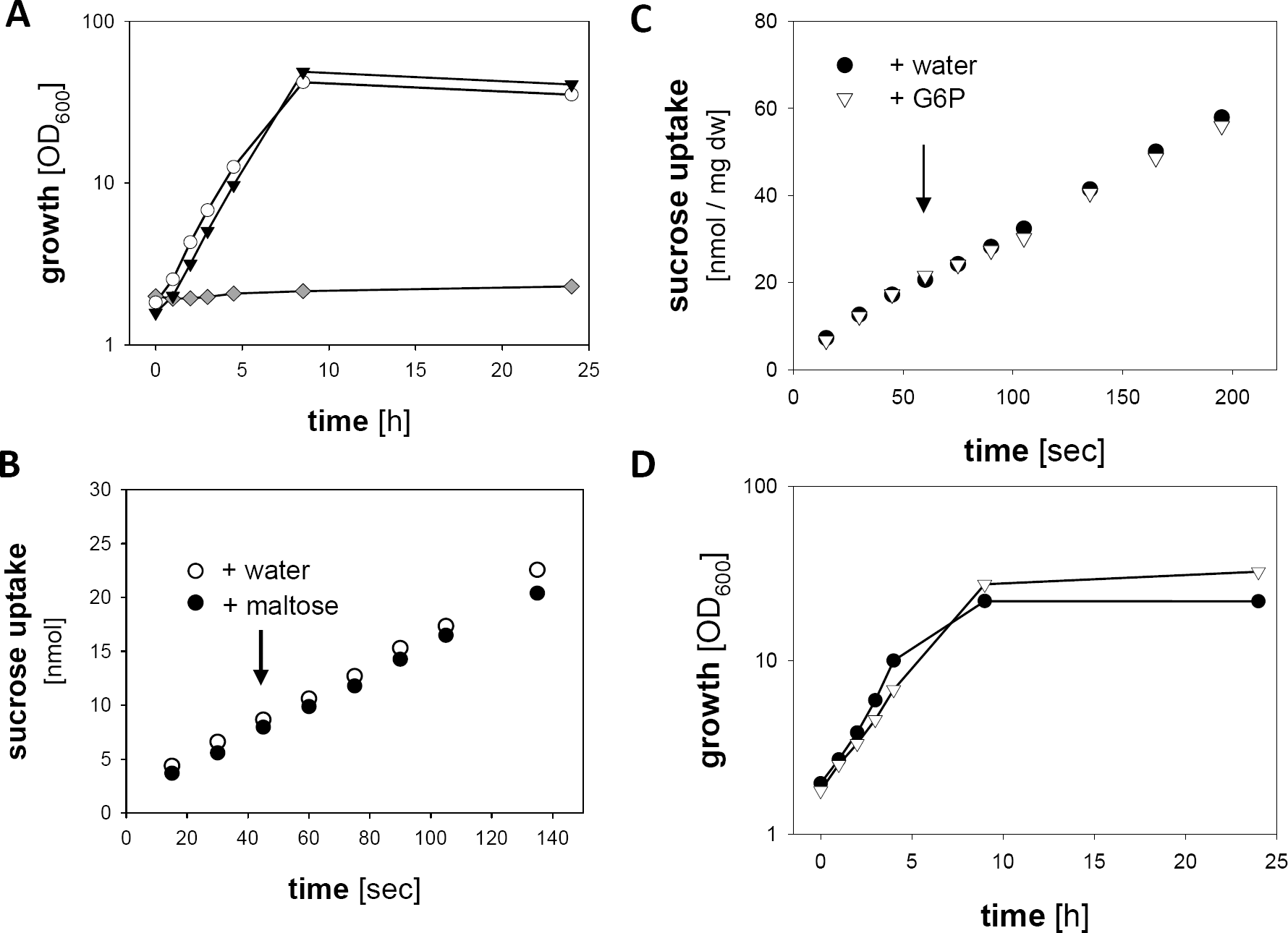
Growth of *C. glutamicum* Δ*pgi* Δ*ptsG* [A] in CgC minimal medium with 2% (w/v) sucrose [black triangles], 1% (w/v) sucrose plus 1 % (w/v) maltose [white circles], or 2% glucose (w/v) [grey diamonds] as carbon sources. Kinetics of [^14^C]-sucrose uptake in response to maltose addition in cells of *C. glutamicum* Δ*pgi* Δ*ptsG* [B] and G6P addition to cells of *C. glutamicum* Δ*pgi* Δ*ptsG* (pEKEx2-*uhpT*) [C]; at the time point indicated by the arrow either 70 μl of 200 mM maltose or G6P solution (white triangles) or 70 μl of water (black circles) were added, one representative experiment of a series of at least three individual experiments is shown. Growth of *C. glutamicum* Δ*pgi* Δ*ptsG* (pEKEx2-*uhpT*) [A] in CgC minimalmedium with 2% (w/v) sucrose [white triangles], 1% (w/v) sucrose plus 1 % (w/v) G6P [black circles]. For growth experiments one representative experiment of a series of three individual experiments is shown.

To further analyse the role of EII^Glc^ for the rapid inhibition of sucrose uptake by intracellular G6P accumulation, additional experiments were performed using the strain *C. glutamicum* Δ*pgi*Δ*ptsG* (pEKEx2-*uhpT*). Neither did addition of unlabelled G6P affect sucrose uptake of the EII^Glc^ deficient strain (Fig. 7C) nor was the growth of *C. glutamicum* Δ*pgi*Δ*ptsG* (pEKEx2-*uhpT*) on sucrose negatively affected by the presence of G6P in the culture broth (Fig. 7D). Growth rates of 0.42 ± 0.02 h^−1^ and 0.38 ± 0.03 h^−1^ were determined for cultivations of *C. glutamicum* Δ*pgi* Δ*ptsG* (pEKEx2-*uhpT*) on sucrose or sucrose plus G6P, respectively. As a control, growth on G6P as sole substrate proceeded in *C. glutamicum* Δ*pgi*Δ*ptsG* (pEKEx2-*uhpT*) with a similar rate of 0.17 ± 0.02 h^−1^) as described above for both *C. glutamicum* (pEKEx2-*uhpT*) and *C. glutamicum* Δ*pgi* (pEKEx2-*uhpT*).

In *E. coli* toxic, intracellular sugar-phosphates, when accumulated to toxic concentrations, are dephosphorylated and then exported (28). Hence, if the accumulating G6P in *C. glutamicum* Δ*pgi* is intracellularly dephosphorylated, and if the resulting glucose is then exported from the cell and eventually reimported by EII^Glc^, a futile cycle would arise leading to consumption of PEP even if the G6P accumulation is not generated by the PTS. PEP consumption would then limit the EII^Suc^ activity albeit the import of the initial substrates itself, G6P via UhpT and maltose via MusEFGK_2_I, does not lead to PEP consumption. In such a scenario, the EII^Glc^-deficient *C. glutamicum* Δ*pgi* Δ*ptsG* strains would also not be affected by intracellular G6P accumulation as they will not consume PEP for glucose reimport. Based on this assumption the EII^Glc^-deficient strains should at least transiently accumulate glucose in the supernatant during cultivation in presence of G6P or maltose. No transient extracellular accumulation of glucose was detected for *C. glutamicum* Δ*pgi*Δ*ptsG* and *C. glutamicum* Δ*pgi*Δ*ptsG* (pEKEx2-*uhpT*) in cultivations with maltose or G6P, respectively. As the reimport of labelled glucose is impossible for the EII^Glc^-deficient strain, export of glucose would result in a decreased rate of label incorporation in these cells. In the course of ^14^C-G6P uptake assays no difference in kinetics for the incorporation of label into cells of *C. glutamicum* Δ*pgi* (pEKEx2-*uhpT*) and *C. glutamicum* Δ*pgi*Δ*ptsG* (pEKEx2-*uhpT*) were observed and thus nearly identical G6P uptake rates of 13.5 ± 2.1 nmol * (min * mg dw)^−1^ and 12.4 ± 0.8 nmol * (min * mg dw)^−1^ were determined [Fig. S11], respectively. In detail, addition of a 100-fold excess (5 mM final concentration) of unlabelled glucose did not cause a reduction of the uptake of ^14^C-labelled G6P in *C. glutamicum* Δ*pgi* (pEKEx2-*uhpT*) [Fig. S11A] and in *C. glutamicum* Δ*pgi*Δ*ptsG* (pEKEx2-*uhpT*) [Fig. S11B]. Based on these results, glucose export and its reimport via EII^Glc^, which would cause PEP consumption, can be excluded to be responsible for the instantaneous inhibition of sucrose uptake via EII^Suc^. We therefore conclude that rapid inhibition of sucrose uptake in *C. glutamicum* Δ*pgi* requires the *ptsG* encoded EII^Glc^, which acts as a sensor for the intracellular accumulation of G6P.

## DISCUSSION

Tight control of substrate uptake is a regulatory prerequisite for bacteria to achieve metabolite homeostasis. Besides its function for uptake and concomitant phosphorylation of substrates the PTS is often a major player in metabolite homeostasis by coordinating substrate uptake when multiple carbon sources are available (3, 4). In detail, the uptake and concomitant phosphorylation of a substrate directly alters the phosphorylation state of PTS components, which than can trigger via protein-protein interaction diverse regulatory cascades supressing the uptake of alternative substrates (6, 38). Here, the glucose-specific EII^Glc^ permease of *C. glutamicum* was shown to cause fast inhibition of the sucrose-specific EII^Suc^ permease upon intracellular accumulation of G6P. This fast inhibition of EII^Suc^ by EII^Glc^ occurred independently of the presence of the substrate of EII^Glc^ as well as of its activity. This control mechanism therefore appears to adapt the activity of EII^Suc^ to intracellular metabolite concentrations and metabolic fluxes. This finding adds a novel aspect to the current understanding of how the PTS activity is controlled, which is usually just regarded to depend on PEP availability and thus coupled to glycolytic flux (39, 40). In *C. glutamicum* excess PEP would be formed in the course of sucrose metabolization [(Fig. 1)]: For uptake and phosphorylation of sucrose and consecutively formed fructose two PEP molecules are required. However, four PEP molecules are formed from sucrose in the course of glycolysis (9, 12). By using the non-PTS substrates Maltose and G6P, which both cause intracellular formation of G6P without requiring PEP consumption for their uptake, we uncoupled substrate uptake and accumulation of intracellular G6P from PEP consumption. Moreover, presence of a futile cycle including dephosphorylation of G6P and reimport of the resulting glucose by EII^Glc^ and thus PEP consumption has been excluded. Therefore, we conclude that G6P accumulation and not PEP-limitation is the primary inducer of the rapid sucrose uptake inhibition in *C. glutamicum* Δ*pgi*. Studies with *E. coli* mutant strains lacking glycolytic enzymes like phosphofructokinase A, phosphoglycerate kinase, glyceraldehyde-3-P dehydrogenase, or enolase cultivated with glucose also showed that the concentration of intermediates upstream of the missing enzyme were increased but the concentration of intermediates downstream of that point, including PEP, remained unchanged when compared to the wild-type (41, 42). Accumulation of G6P and F6P were shown to trigger the SgrS-dependent degradation of *ptsG* in *E. coli* and the concomitant inhibition of glucose uptake (43). Monitoring metabolite levels both at the entrance and exit of glycolysis, probably allows the cell to coordinate more precisely the co-utilisation of different substrates as opposed to a response to a single downstream metabolite. This dual control of the PTS prohibits in *E. coli* the detrimental effects of sugar-phosphate accumulation when PEP becomes available, *e.g* caused by the presence of pyruvate in the culture broth (44). In general, inhibition of the EII permease by glycolytic intermediates is a plausible way to adapt PTS activity to the metabolic capacities. In *E. coli* activity of the EII-permeases EII^Glc^ and EII^Man^ for glucose uptake is inhibited by accumulation of G6P and other glycolytic intermediates (45-47), which contributes to the fast adjustment of metabolite concentrations upon substrate changes (48). However, in contrast to EII^Glc^ and EII^Man^ of *E. coli*, which are directly responding to accumulation of G6P (46), in *C. glutamicm* the control of the activity of EII^Suc^ in response to G6P accumulation is indirect and requires the presence of EII^Glc^. Albeit the molecular mechanism underlying the signalling process remains elusive, this ability of EII^Glc^ in *C. glutamicum* to sense intracellular G6P levels and to transduce a specific signal to EII^Suc^ represents a novel functionality of an EII-permease. The *ptsG* encoded EII^Glc^ as well as the *ptsS* encoded EII^Suc^ in *C. glutamicum* each consist of fused EIIA, EIIC and EIIB domains (12). This organisation of the EII permeases in *C.glutamicum* is different from the situation in *E. coli*, in which the EIIA of the PTS is not covalently linked to the permease domain EIIC (39). Thus, EIIA in *E. coli* can easily exert multiple regulatory functions via direct binding to regulatory sites of different substrate importers, *e.g.* the LacY permease (49, 50) and the maltose ABC transporter (51, 52). Based on the different architecture of the *C. glutamicum* EII^Glc^ and EII^Suc^ a different mode of interaction between domains of two EII permeases is necessary to bring along the observed fast inhibition of sucrose uptake in response to G6P accumulation. The mechanistic details for this interaction remain to be investigated.

Transcription of the *ptsS* gene, encoding for EII^Suc^, also was found to be reduced in *C. glutamicum* Δ*pgi* in response to glucose addition, which is a regulatory reaction not observed for *C. glutamicum* WT and *C. glutamicum* Δ*pgi* Δ*sugR*. The DeoR-type transcriptional repressor SugR is a global regulator in *C. glutamicum* for the transcriptional control of genes for sugar uptake and their metabolization via glycolysis (33, 34, 53, 54). SugR binding sites are located within the promoter regions of genes for PTS components *e.g*. within the promoter of *ptsS* (55). Fructose-1-P has been identified as a potential effector of SugR, triggering *in vitro* the release of SugR from its target promoters (34). Fructose-1-P is formed in *C. glutamicum* in the course of sucrose metabolization exclusively via the *ptsF*-encoded, fructose specific PTS-permease EII^Fru^, which is required for the re-uptake and concomitant phosphorylation of the fructose formed by ScrB (12, 32, 56) (Fig. 1). Based on this metabolic pathway for sucrose utilization a rapid inhibition of sucrose uptake upon addition of glucose will limit formation of fructose-1-P via EII^Fru^, as less fructose is intracellularly formed by ScrB from sucrose-6-P and exported. Thus, in an inducer exclusion-like manner, the fast inhibition of sucrose uptake by glucose addition might lead to decreased levels of the SugR effector molecule fructose-1-P, which probably explains the SugR-dependent repression of *ptsS* observed upon prolonged incubation of *C. glutamicum* Δ*pgi* in presence of sucrose plus glucose.

The initial inhibition of EII^Suc^ by EII^Glc^ in response to accumulation of the intermediate G6P, is probably a first insight into the complex architecture of regulatory mechanisms underlying the ability of *C. glutamcium* to adapt substrate uptake perfectly to its needs, a typical trait of this bacterium (14). This leads to the question of whether the described EII^Glc^-dependent rapid uptake inhibition is a general feedback response mechanism to accumulation of G6P in *C. glutamicum* thus influencing the activity of other sugar transporters besides EII^Suc^. Indeed recent observations in our lab concerning the control of maltose utilization by the PTS in *C. glutamicum* (57) indicate that the PTS and its components might have a more general, hitherto often neglected role in the control of carbohydrate metabolism in this organism. Taken together, we show that besides its effect on transcriptional regulation, *C. glutamicum* possesses a novel, immediate, and PTS-dependent way to control and coordinate both uptake and metabolization of multiple substrates by monitoring of their metabolic levels in the cell. This offers new insights and interesting concepts for a further rational engineering of this industrially important production organism. The reported finding of a PEP-independent, tight control of PTS-mediated sugar uptake in response to the accumulation of the initial glycolysis intermediate G6P in *C. glutamicum*, in addition to previous findings for G6P and F6P dependent control of PTS activity in *E. coli*, further exemplifies a probably general strategy of bacteria for the coordination of sugar uptake and central metabolism.

## MATERIALS AND METHODS

### Bacterial strains, media and culture conditions

Bacteria and plasmids used in this study are listed in Table 1. *E. coli* and all pre-cultures of *C. glutamicum* were grown aerobically in TY complex medium (58) at 37 °C and 30 °C, respectively, as 50-ml cultures in 500-ml baffled Erlenmeyer flasks on a rotary shaker at 120 rpm. For the main cultures of *C. glutamicum*, cells of an overnight pre-culture were washed twice with 0.9% (w/v) NaCl and then inoculated into CGC minimal medium (59) containing the carbon sources as indicated in the results section. When appropriate, kanamycin (50 μg ml^−1^) and/or isopropyl-β-D-thiogalactopyranoside (IPTG, 250 μM) were added to the media. Growth of *E. coli* and of *C. glutamicum* was followed by measuring the optical density at 600 nm (OD_600_) in an Ultrospec 2000 spectrophotometer (GE Healthcare).

**Table 1:**
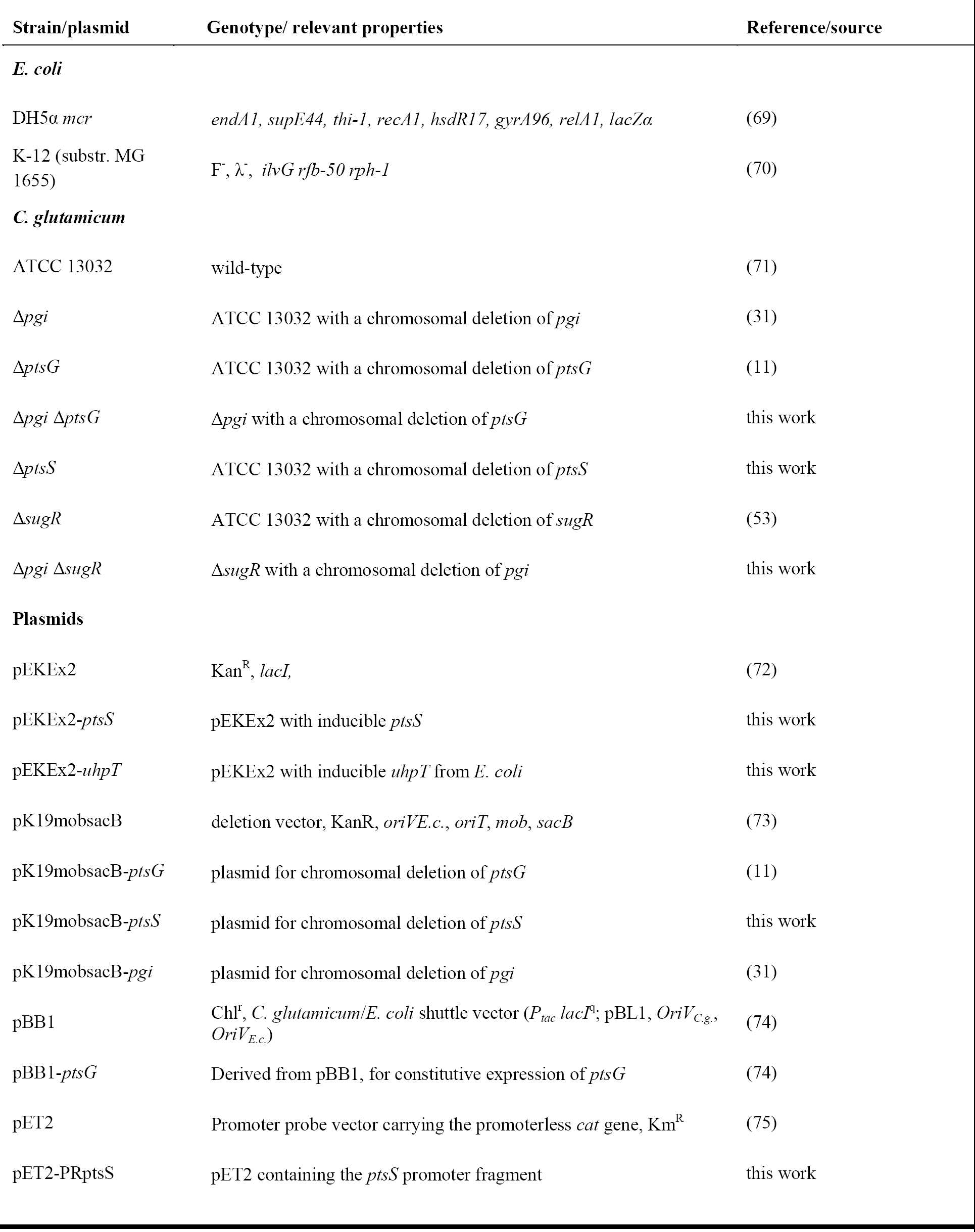
Bacterial strains and plasmids used during this work

### DNA isolation, transfer and manipulations

Standard procedures were employed for plasmid isolation, cloning, and transformation of *E. coli* DH5α, as well as for electrophoresis (58). *C. glutamicum* chromosomal DNA was isolated according to (60). Transformation of *C. glutamicum* was performed by electroporation using the methods of (61), the recombinant strains were selected on LB-BHIS agar plates containing kanamycin (25 μg ml^−1^). Electroporation of *E. coli* was performed according to the method of (62). All enzymes used were obtained from New England Biolabs and used according to the instructions of the manufacturer. PCR experiments were performed in a TProfessional thermocycler (Biometra). Desoxynucleoside triphosphates were obtained from Bio-Budget, oligonucleotides (primers) from Eurofins MWG Operon. Cycling times and temperatures were chosen according to fragment length and primer constitution. PCR products were separated on agarose gels and purified using the Nucleospin extract II kit (Macherey& Nagel).

### Construction of expression vectors

For IPTG-inducible overexpression vector pEKEx2 was used. Genes were amplified via PCR from genomic DNA of *C. glutamicum* ATCC 13032 using the oligonucleotide primers listed in Table S1. The resulting PCR products were introduced into the cloning vector pJET1.2 (MBI Fermentas) according to the manufacturer’s instructions. Primer-attached restriction sites of the PCR products (indicated in Table S1) were used to excise the inserts, the resulting fragments were ligated into the plasmid pEKEx2 (digested with appropriate enzymes) and transformed into *E. coli* DH5α. The resulting plasmids were isolated and the nucleotide sequences controlled by sequencing (GATC Biotech).

### Construction of the deletion mutants

The in-frame deletion of *ptsS* in *C. glutamicum* was performed via a two-step homologous recombination procedure as described previously (Niebisch and Bott, 2001). In detail, the flanking up- and downstream regions of *ptsS* were amplified using primer pairs ptsS_A and ptsS_B and ptsS_C and ptsS_D and the product of the following crossover PCR was cloned as a BamHI-XbaI cut fragment into the pK19*mobsacB* vector (Schäfer *et al*., 1994). Transfer of the resulting plasmid, pK19mobsacB-*ptsS*, into *C. glutamicum* by electroporation and screening for the correct mutants was performed as described previously for the deletion of *ptsG* (11). The deletion of *ptsS* was verified using the primers ptsS_chkA and ptsS_chkB, resulting in 3926 bp fragment for the WT and a 1966 bp fragment for the deletion mutant. The in-frame deletion of *ptsG* in *C. glutamicum* Δ*pgi* was performed as described previously using plasmid pK19*mobsacB*-*ptsG* (11), and verified by PCR using the primers ptsG_chkA and ptsG_chkB. In-frame deletion of *pgi* in *C. glutamicum* Δ*sugR* was performed as described previously using the plasmid pK19*mobsacB*Δ*pgi* and verified by PCR using the primers pgi_chkA and pgi_chkB, as recently described (Lindner *et al*., 2013).

### Analysis of cytoplasmic and extracellular glucose-6-P concentrations

Rapid sampling, inactivation of metabolism and separation of intracellular and extracellular fluids for enzymatic analyses of intracellular G6P concentrations cells was achieved by silicone oil centrifugation as recently described (63). The samples were neutralized with 25 μl of 1 M KOH, 5 M triethanolamine, potassium perchlorate thereby generated was precipitated by incubation for 30 min at 4 °C followed by centrifugation (5 min, 20000 ***g***, 4 °C). The supernatant was transferred to a new vial and used for the enzymatic determination of G6P. The assay contained the sample and 2mM NADP in a final volume of 240 μl, adjusted with G6-buffer (0.15 m TEA, 10 mM MgSO_4_, 0.1 M KCl). The measurement was started with the addition of 10 μl glc-6-P dehydrogenase (0.1 U). The absorbance of the samples was measured photometrically in Ultrospec 2000 spectrophotometer at a wavelength of 340 nm at constant temperature of 37 °C. For the measurement of extracellular G6P concentrations when G6P was used as a carbon source, the same method was used, however, by measuring the supernatants instead of cell extracts obtained after the silicon oil centrifugation.

### Chloramphenicol acetyl-CoA transferase (CAT) assay

To study *ptsS* promoter activity a transcriptional fusion of the *ptsS* promoter to the promoterless *cat* gene within the reporter plasmid pET2 was constructed. Therefore, the *ptsS* promoter fragment was amplified by PCR using primers PRptsS-for and PRptsS-rev (Table 3), the resulting 322 bp PCR product (covering the region 78 bp upstream to 243 bp downstream of the *ptsS* transcriptional start codon) was cloned into the multiple cloning site in front of the *cat* gene in pET2 using restriction sites indicated in Table 3 resulting in pET2-PR*ptsS*. To determine CAT activity, *C. glutamicum* cells were harvested during the exponential growth phase, washed twice in 0.1 M Tris/HCl, pH 7.8, and resuspended in the same buffer containing 10 mM MgCl_2_ and 1 mM EDTA. The specific CAT activity was determined as described by Schreiner *et al*. (64). Protein concentrations were determined using the Roti-Nanoquant kit (Roth) with bovine serum albumin as the standard.

### RNA techniques

Isolation of total RNA from *C. glutamicum* cells was performed using the Nucleospin® RNAII kit (Macherey& Nagel) as described by (65). For Northern (RNA) hybridisation, digoxigenin (DIG)-11-dUTP-labeled gene specific antisense RNA probes were prepared from PCR products (generated with oligonucleotides listed in Table S1) carrying the T7 promoter by *in vitro* transcription (1h, 37° C) using T7 RNA polymerase (MBI Fermentas). Slot blot experiments, detection, and signal quantification were performed as described previously (31, 66).

### Uptake studies

Sugar uptake studies were performed essentially as recently described for the uptake of maltose (11). In detail, *C. glutamicum* cells were grown in the media indicated in the text to mid-exponential growth phase, harvested by centrifugation, washed three times with ice cold CgC medium, suspended to an OD_600_ of 2 with CgC-medium, and stored on ice until the measurement. Before the transport assay, cells were tempered for 3 min at 30 °C; the reaction was started by addition of ^14^C (U) radioactively labelled maltose, glucose-6-P or sucrose (American Radiolabeled Chemicals Inc, Saint Louis, USA). At given time intervals (resolution of 15 sec), 200 μl samples were filtered through glass fibre filters (Typ F; Millipore) and washed twice with 2.5 ml of 100 mM LiCl. Radioactivity of the samples was determined using scintillation fluid (Rotiszinth; Roth) and a scintillation counter (LS 6500; Beckmann).

If not indicated otherwise, the substances tested as inhibitors in the rapid uptake studies were added to the setup 45 sec or 60 sec (indicated in figure legends) after the measurement was started by addition of the ^14^C-labelled substrate and the potential inhibitors were used in at least 10-fold excess compared to the concentration of the labelled substrate (concentrations are provided in the figure legends). For transport experiments performed in absence of transcriptional or translational regulatory processes the cells were treated before the addition of the labelled substrate with rifampicin or chloramphenicol, respectively, as described by (67, 68).

## Abbreviations

cdm: cell dry mass
WT: wild-type

## Acknowledgments

We thank Bernhard Eikmanns for continuous support and the German Ministry of Education and Research for financial funding in the frame of the e:Bio initiative (contract no. 031A302D). We are very grateful to Volker Wendisch for the generous gift of the strain *C. glutamicum* Δ*sugR*.

